# A scalable computational framework for predicting gene expression from candidate cis-regulatory elements

**DOI:** 10.1101/2025.07.21.666040

**Authors:** Qinhu Zhang, Siguo Wang, Zhipeng Li, Wenzheng Bao, Wenjian Liu, De-Shuang Huang

## Abstract

Deciphering the relationships between cis-regulatory elements (CREs) and target gene expression has been a long-standing unsolved problem in molecular biology, and the dynamics of CREs in different cell types make this problem more challenging. To address this challenge, we propose a <u>**sc**</u>alable computational framework for <u>**p**</u>redicting <u>**g**</u>ene <u>**e**</u>xpression (ScPGE) from discrete candidate CREs (cCREs). ScPGE assembles DNA sequences, transcription factor (TF) binding scores, and epigenomic tracks from discrete cCREs into 3-dimensional tensors, and then models the relationships between cCREs and genes by combining convolutional neural network with transformer. Compared with current state-of-the-art models, ScPGE exhibits superior performance in predicting gene expression and yields higher accuracy in identifying active enhancer-gene interactions through attention mechanisms. By comprehensively analyzing ScPGE’s predictions, we find a pattern in true positives (TPs) that the regulatory effect of cCREs on genes decreases with distance. Inspired by the pattern, we design two methods to enhance the ability to capture distal cCRE-gene interactions by incorporating chromatin loops into the ScPGE model. Furthermore, ScPGE accurately discovers some crucial TF motifs within prioritized cCREs and reveals the different regulatory types of these cCREs.

## 1. Introduction

Cell-type-specific gene expression patterns are primarily determined through complex interactions between cisregulatory elements (CREs) and transcriptional factors, playing a crucial role in differentiation and development(Furlong and Levine 2018). Mutations in CREs contribute to various diseases by disrupting the regular expression of their target genes(Nasser et al. 2021). Decoding how CREs regulate gene expression may reveal the mechanisms of gene regulation and provide insights into the origins of human diseases. However, epigenomic data, such as chromatin accessibility and histone modifications, enhance the dynamic characteristics of CREs to gene regulation, making it more challenging to decipher the relationships between CREs and gene expression.

High-throughput experimental techniques, such as Hi-C(Rao et al. 2014), ChIA-PET(Tang et al. 2015), and HiChIP(Mumbach et al. 2016), have been developed for identifying physical CRE-gene interactions in a genome-wide fashion, but they just measure physical proximity between CREs and genes instead of direct regulatory impact. Recently, systematic evaluation of the impact of CREs on gene expression has become possible with CRISPR perturbations(Fulco et al. 2019), but only a small subset of CREs can be evaluated in the genome. Meanwhile, the evaluation is restricted to a small number of cell types. Due to its high cost, it is difficult to apply on a large scale.

Quantitative models have been proposed recently for modelling the relationships between candidate CREs (cCREs) and gene expression. Early models mainly focus on forming binary classification tasks to predict physical cCRE-gene interactions(Li et al. 2019; Tang et al. 2020; Cao et al. 2021) rather than directly predicting gene expression regulated by cCREs. Meanwhile, their performance and generalization ability are always subject to the number of real cCRE-gene interactions, the varying number of cCREs for target genes, and the complex nature of cCRE-gene regulation(Schoenfelder and Fraser 2019). Interestingly, recent studies on predicting gene expression from DNA sequences have shown that transformer-based algorithms implicitly incorporate modelling cCRE-gene interactions and significantly improve the performance of gene expression prediction(Avsec et al. 2021; Li et al. 2023; Lin et al. 2024). Their special attention mechanisms can effectively capture long-range cCRE-gene interactions, offering a significant advantage over convolutional neural networks (CNN), which focus on local interactions. Enformer(Avsec et al. 2021), a representative transformer-based model, excels in predicting gene expression, chromatin states, and variant effects. However, since this method only uses sequences, it cannot recognize cell-type-specific cCREs, signifying that its adaptability to unseen data from new cell types is limited(Sasse et al. 2023). Moreover, training a new model from scratch using this method requires a substantial amount of computing resources, posing a significant challenge for researchers with limited computing capabilities. To address these challenges, several models of modest size have been proposed to improve the performance of gene expression prediction by utilizing extra data. For example, GraphReg(Karbalayghareh et al. 2022) introduced a graph attention network that integrates chromatin contact data (Hi-C) to predict gene expression. CREaTor(Li et al. 2023) presented a hierarchical deep learning model based on the self-attention mechanism, which utilizes cCREs in open chromatin regions together with ChIP-seqs of transcription factors and histone modifications to predict the expression level of target genes. EPInformer(Lin et al. 2024) introduced an efficient deep-learning framework based on the transformer architecture to predict gene expression by integrating DNA sequences, epigenomic signals, and chromatin contact data in stages. Although these models achieve remarkable predictive performance, their effectiveness is constrained by the scarce availability of extra data across different cell types or by the requirement of elaborately preparing data.

In this study, we introduce a scalable computational framework for predicting gene expression (ScPGE) from discrete cCREs identified by the Encyclopedia of DNA Elements (ENCODE). ScPGE first assembles DNA sequences, TF binding scores, and epigenomic tracks from discrete cCREs into three 3-dimensional tensors, respectively, and then combines CNN with transformer to predict gene expression levels. In the ScPGE’s architecture, CNN serves as a feature extractor to learn the local sequence or chromatin features, while transformer serves as a relationship extractor to learn the relationships between genes and cCREs, as well as cCREs and cCREs. Notably, ScPGE demonstrates its flexible scalability from three aspects: (i) scalability of epigenomic tracks, enabling the use of any number of epigenomic tracks if available; (ii) scalability of cCREs, allowing for the incorporation of any number of cCREs if reasonable; (iii) scalability of model structure, adjusting the layers of CNN and transformer according to the sequence length. Moreover, ScPGE greatly reduces computational demands for modelling by using discrete cCREs instead of the entire genomic region flanking target genes, facilitating rapid training and deployment for new cell types. Experimental results confirm the superiority of ScPGE over existing models, showing that ScPGE can predict RNA-seq or CAGE-seq gene expression accurately by modelling the relationships between cCREs and target genes effectively. Further analysis of predictions reveals different regulatory patterns, finding a pattern in true positives (TPs) that the regulatory effect of cCREs on genes decreases with distance. With this pattern, ScPGE can perform cross-cell or cross-species gene expression prediction, implying its potential for application to unseen cell types or species. Moreover, inspired by this pattern, we design two methods to enhance the ability to capture distal cCRE-gene interactions by incorporating chromatin loops into the ScPGE model. We believe that the concept of ScPGE can provide valuable insights for studying gene regulation under resource-limited conditions.

## 2. Results

### 2.1 Overview of ScPGE

ScPGE is a deep learning framework, composed of a feature-learning module, an interaction-learning module, and a prediction module, which predicts cell-type-specific gene expression levels by integrating DNA sequences, TF binding scores, and epigenomic signals from discrete cCREs. As shown in Fig.1, ScPGE first converts DNA sequences, TF binding scores, and epigenomic signals into three 3-dimensional tensors, respectively. Secondly, ScPGE takes the three tensors as input and utilizes multiple *2D* convolutional blocks to learn multi-modal features of cCREs in parallel, as well as transformer-based blocks to model the relationships between genes and cCREs. Finally, ScPGE extracts the features of cCRE-gene pairs and predicts gene expression levels through the prediction module. Importantly, given that chromatin loops could provide useful information for guiding ScPGE to capture the relationships between genes and cCREs, two approaches are developed to incorporate chromatin loops into the ScPGE model by either increasing the attention weights of chromatin interactions directly or introducing a KL Divergence loss indirectly (**Methods**).

**Fig.1.**
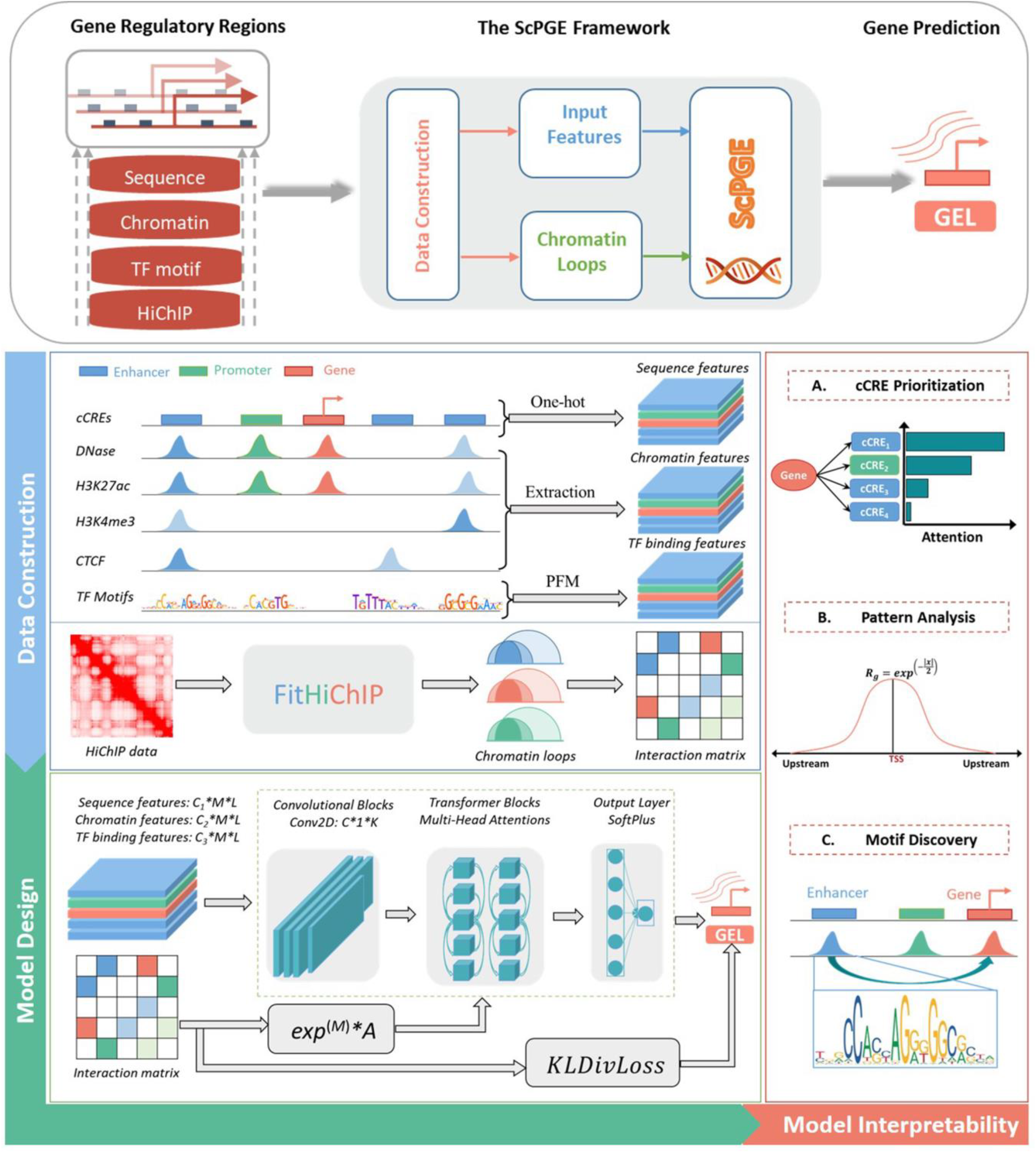
The framework of ScPGE. The framework predicts gene expression by integrating sequence features, chromatin features, TF motif features, and HiChip interactions of discrete candidate cCREs on both sides of genes. During the stage of data construction, we transformed sequence features, chromatin features, and TF motif features into 3-dimensional tensors, and converted chromatin loops identified by FitHiChIP into interaction matrices. During the stage of model design, we combined CNN and transformer to predict gene expression, where CNN is used to learn sequence and chromatin features while transformer is used to learn the relationships between genes and cCREs. Meanwhile, we designed two ways to improve the performance of ScPGE by incorporating chromatin loops. In the stage of model interpretability, we performed cCRE prioritization, pattern analysis, and motif discovery.

ScPGE was trained to minimize the discrepancy between predicted and observed gene expression levels, as measured by RNA-seq or CAGE-seq, using multiple cell types. For each cell type, all coding genes from chromosome 16 were used for validation, all coding genes from chromosomes 8 and 9 were used for testing, and all coding genes from the remaining chromosomes, except chromosome Y, were used for training.

### 2.2 ScPGE accurately predicts gene expression across multiple cell types

In this section, we used three state-of-the-art models such as Enformer(Avsec et al. 2021), CREaTor(Li et al. 2023), and EPInformer(Lin et al. 2024) to compare with ScPGE and quantified their performance by the Pearson’s correlation coefficient (PCC) and mean absolute error (MAE) between predicted and observed gene expression levels in a variety of cell types.

To assess the performance of ScPGE in predicting RNA-seq gene expression levels, we trained ScPGE, CREaTor, and EPInformer by utilizing RNA-seq datasets from 19 human cell types. As shown in Fig.2a, ScPGE consistently outperforms both CREaTor and EPInformer, achieving significant performance improvements. Specifically, ScPGE surpasses CREaTor’s performance by 2.5% and 27% in PCC and MAE, respectively, and exceeds EPInformer’s performance by 1.3% and 1% in PCC and MAE, respectively.

**Fig.2.**
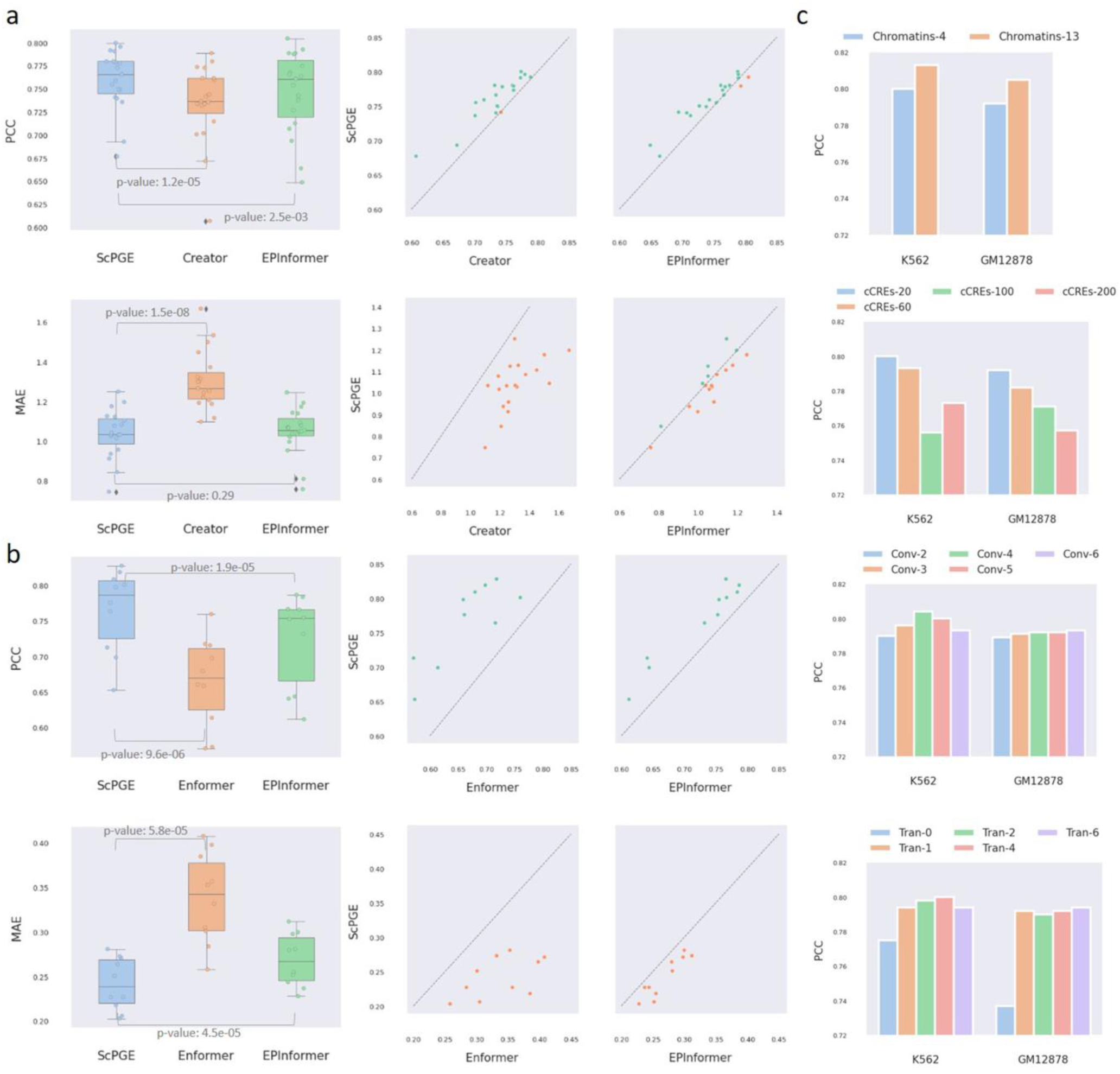
Overall performance of ScPGE. (a) The performance of ScPGE in predicting RNA-seq gene expression levels, which was measured by PCC and MAE. (b) The performance of ScPGE in predicting CAGE-seq gene expression levels, which was measured by PCC and MAE. The *p*-value is calculated using student’s T-test. (c) The performance (PCC) of ScPGE’s scalability by modifying the number of epigenomic tracks, the number of cCREs, and the structure of ScPGE.

To evaluate the performance of ScPGE in predicting CAGE-seq gene expression levels, we trained ScPGE and EPInformer by utilizing CAGE-seq datasets from 10 human cell types available in Enformer and compared them with Enformer’s pre-trained models. As illustrated in Fig.2b, ScPGE significantly outperforms both Enformer and EPInformer across all datasets. ScPGE shows improvements of 10% in PCC and 9.6% in MAE compared to Enformer, and improvements of 4.2% in PCC and 2.7% in MAE compared to EPInformer.

In summary, the above results obtained from RNA-seq and CAGE-seq datasets indicate that ScPGE can accurately predict gene expression across various cell types, highlighting its effectiveness in predicting gene expression.

To demonstrate the flexible scalability of ScPGE, we performed a series of experiments by modifying the number of epigenomic tracks, the number of cCREs, and the structure of ScPGE. (i) Scalability of epigenomic tracks (Fig.2c): we find that ScPGE using 13 epigenomic tracks performs better than ScPGE using 4 default tracks, suggesting that a greater number of epigenomic tracks provides more valuable information. (ii) Scalability of cCREs (Fig.2d): we observe a decrease in ScPGE’s performance as the number of cCREs increases, implying that more cCREs may introduce noise rather than useful information. (iii) Scalability of model structure (Figs.2e-f): we notice that the lack of transformer (Tran-0) significantly reduces the predictive performance of ScPGE, while other cases had little impact on its performance, highlighting the crucial role of transformer in modelling the relationships between genes and cCREs. Consequently, ScPGE exhibits flexible scalability by accommodating any number of epigenomic tracks and cCREs, as well as allowing for easy adjustment of the model architecture, which makes it particularly well-suited for gene expression prediction in varying contexts, including situations where data availability is uncertain.

### 2.3 ScPGE successfully prioritizes cell-type-specific cCREs

Linking candidate enhancers to their target genes through high-throughput experiments remains a critical challenge in terms of time and cost. In this scenario, accurate prioritization of candidate enhancer-gene interactions by computational methods becomes increasingly important. To evaluate ScPGE’s accuracy in capturing true enhancer-gene interactions, we focus on genes with experimentally validated enhancers. These enhancer-gene interactions were collected from three public literatures, consisting of positive (677) and negative (2239) interactions, respectively, and categorized into four groups based on distance.

After the K562-specific ScPGE model was trained, the attention scores from self-attention layers were used to prioritize all candidate enhancer-gene interactions (Fig.3a). Additionally, *in-silico* perturbation based on ScPGE was performed to prioritize all candidate enhancer-gene interactions by their relative changes. As measured by the PRAUC metric (Fig.3b), the performance of ScPGE is comparable to EPInformer and superior to Enformer and CREaTor at various distances, demonstrating the effectiveness of ScPGE. Furthermore, the performance of all methods decreases with increasing distance, suggesting that they perform poorly in predicting distal enhancer-gene interactions.

**Fig.3.**
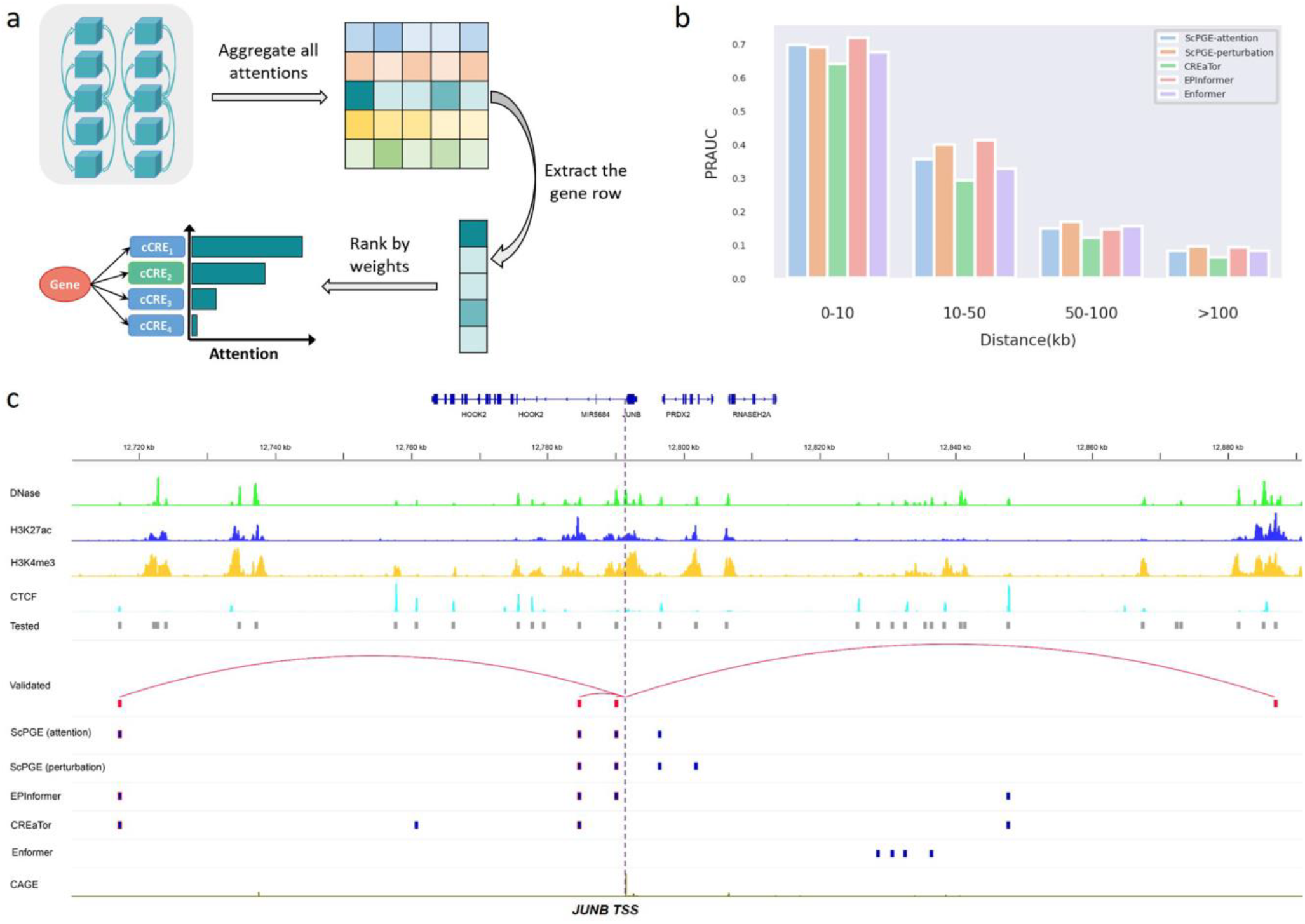
ScPGE prioritizes cell-type-specific cCREs. (a) Schematic overview of prioritizing active cCREs by ScPGE’s attentions. (b) The performance (PRAUC) of relevant methods in classifying cCRE-gene pairs at different distance groups. (c) Visualization of active cCREs of the JUNB gene prioritized by ScPGE (attention), ScPGE (perturbation), EPIformer, CREaTor, and Enformer, where validated cCREs are represented by red squares and correctly identified cCREs are represented by red boxes.

To illustrate ScPGE’s capability in detecting active enhancers for a given locus, we focused on two important genes (JUNB(Chen et al. 2022) and KLF1(Myers et al. 2025)) in the human hematopoietic system. We selected all candidate enhancers surrounding the two genes within 100kb as test targets. These enhancers (grey boxes) were evaluated via CRISPRi-FlowFISH experiments, and only a few were found to have a significant effect on gene expression. To classify candidate enhancers as active or inactive, we prioritized all enhancers and selected top-*k* enhancers, where *k* is the number of validated enhancers, to calculate the true positive rate (TPR). As shown in

Fig.3c, there are four validated enhancers (red boxes) surrounding the JUNB gene, including two proximal enhancers and two distal enhancers. We find that both ScPGE (attention) and EPInformer discover three validated enhancers, achieving the highest TPR of 75%, and outperform other methods. Additionally, we find that ScPGE (attention) tends to capture distal enhancers, while ScPGE (perturbation) tends to capture proximal enhancers. Therefore, they complement each other and can be combined to capture active enhancers. As shown in Supplementary Fig.3, there are six validated enhancers (red boxes) surrounding the KLF1 gene, including three proximal enhancers and three distal enhancers. We can observe that ScPGE (attention + perturbation) discovers all validated enhancers, outperforming other methods in capturing active enhancers.

### 2.4 Categorization of ScPGE’s predictions shows distinct patterns

Although several computational methods for predicting gene expression have been proposed, there is little in-depth analysis of their predictions. Such analysis is essential for identifying distinct patterns from predictions and providing feedback for optimizing models. To achieve this, we categorized ScPGE’s predictions into true positives (TPs), false positives (FPs), true negatives (TNs), and false negatives (FNs) (**Methods**).

Through ablation experiments (Supplementary Fig.4), we observed that chromatin signals were the most significant factor affecting the prediction performance. To uncover distinct patterns from the four types of predictions, we performed a visual analysis of the chromatin signal distributions of cCREs surrounding the transcription start sites (TSS) of genes. For TPs (Fig.4a), we discover a pattern where the distribution of four chromatin signals follows a normal distribution, that is, the intensity of the chromatin signals decreases with distance. In contrast, FPs exhibit a similar normal distribution (Supplementary Fig.5a), which explains why negative genes were mistakenly predicted as positive genes. For TNs (Supplementary Fig.5b), we observe a linear distribution of the four chromatin signals, with DNase and H3K27ac signals close to zero. Similarly, FNs also show a linear distribution of the chromatin signals (Supplementary Fig.5c), particularly with DNase signals close to zero, suggesting that DNase plays a key role in mispredicting positive genes as negative genes. Notably, these patterns indicate that most of coding genes are primarily regulated by their nearby cCREs.

**Fig.4.**
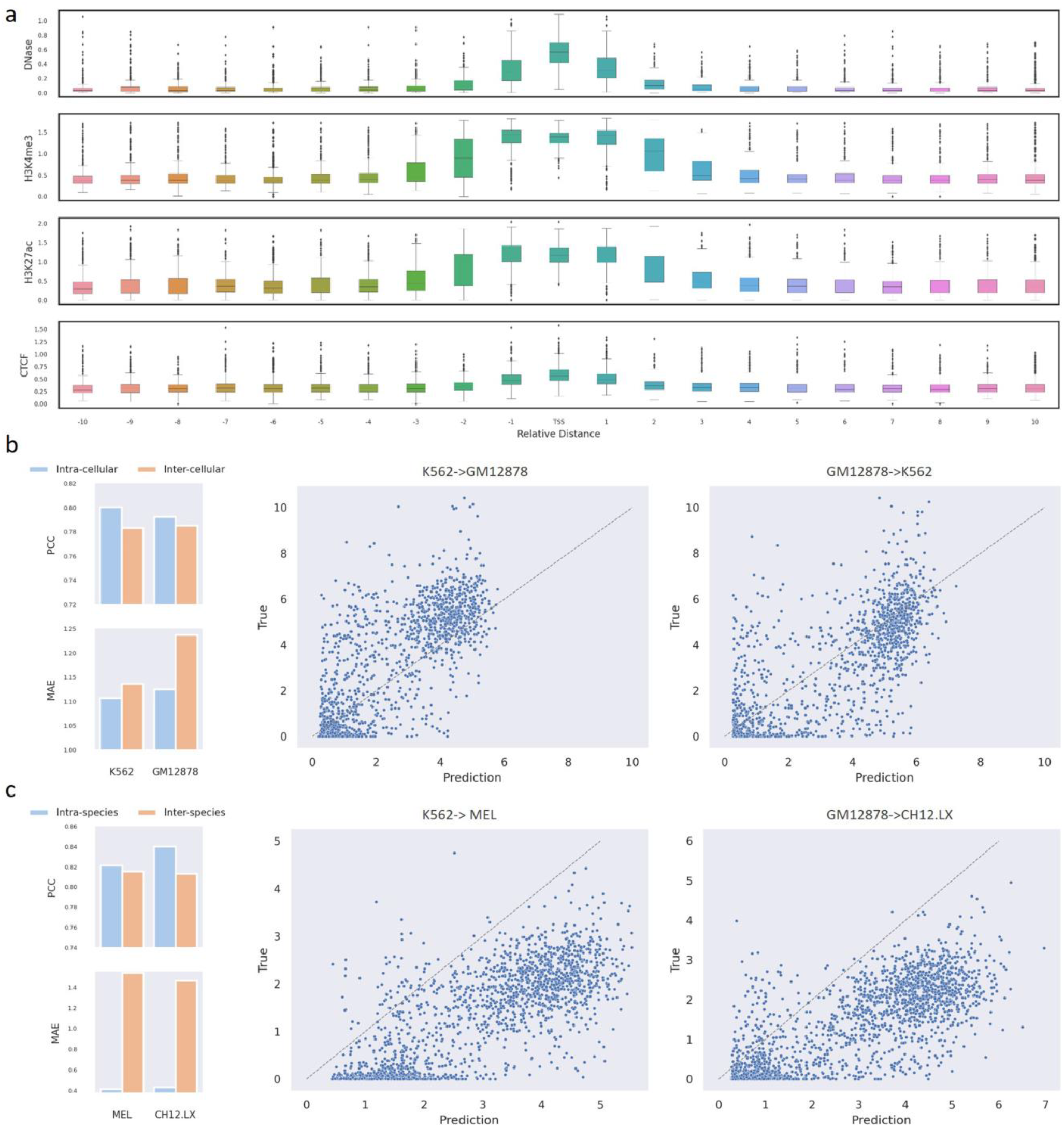
ScPGE discovers different patterns by analyzing predictions. (a) A pattern found in true positives (TPs) that the regulatory effect of cCREs on target genes decreases with distance. (b) The cross-cell type predictive performance of ScPGE, where K562->GM12878 indicates using a model trained on K562 data to predict GM12878 data. (c) The cross-species predictive performance of ScPGE, where K562->MEL indicates using a model trained on K562 data (Human) to predict MEL data (Mouse).

Due to the generalization of the patterns across cell or species (Supplementary Fig.6), we applied ScPGE to perform cross-cell or cross-species gene expression prediction. In the cross-cell setting, we utilized the ScPGE model trained on the GM12878 cell line to predict gene expression levels in the K562 cell line and then vice versa. As shown in Fig.4b and Supplementary Fig.7a, the performance of the inter-cellular prediction is very close to the intra-cellular prediction, with only a slight decrease, demonstrating that ScPGE can effectively generalize the patterns to perform cross-cell gene expression prediction accurately. In the cross-species setting, we utilized the ScPGE models trained on the GM12878 and K562 cell lines to predict gene expression levels in the CH12.LX and MEL cell lines, respectively. As illustrated in Fig.4c and Supplementary Fig.7b, although the PCC of the inter-species prediction is comparable to the intra-species prediction, the MAE of the inter-species prediction is significantly higher than that of the intra-species prediction. This suggests that although ScPGE can capture the general distribution of gene expression, it struggles to accurately predict absolute expression levels due to the strong heterogeneity between different species.

### 2.5 Chromatin loops facilitate the capture of cCRE-gene interactions

In this study, we have shown an issue that the performance of ScPGE decreases as the number of cCREs increases. To alleviate this issue, we designed two methods to incorporate chromatin loops derived from HiChIP into the ScPGE model (**Methods**), enhancing its ability to capture distal cCRE-gene interactions. In the direct method, we directly put chromatin loops into the self-attention layer of ScPGE, aiming to increase the attention weights of validated cCRE-gene interactions. For simplicity, we refer to it as ScPGE-LP. Inspired by the pattern found in TPs that the regulatory effect of cCREs on target genes would decrease with distance, we first introduced an exponential decay function *exp*^−|*x*|/2^ into chromatin loops, aiming to alleviate the sparsity of chromatin loops. Then, we added a KL Divergence loss between chromatin loops and attention weights to the training loss, with the goal of aligning their distributions. For simplicity, we refer to this method as ScPGE-KL.

To evaluate the effectiveness of ScPGE-LP and ScPGE-KL, we trained both models from scratch using 5kb resolution chromatin loops from the K562 and GM12878 cell lines, which represent physical interactions between cCREs and genes. As shown in Figs.5a-b, the performance of ScPGE-LP and ScPGE-KL is superior to that of ScPGE, particularly when utilizing a higher number of cCREs (e.g., cCRE-100, cCRE-200), demonstrating that incorporation of cCRE-gene interactions can improve the performance of predicting gene expression. In addition, ScPGE-KL outperformed ScPGE-LP when a smaller number of cCREs were used (e.g., cCRE-20), suggesting that the regulatory pattern of exponential decay enhances the model’s ability to identify proximal cCRE-gene interactions. In contrast, ScPGE-LP excelled over ScPGE-KL when a higher number of cCREs were employed (e.g., cCRE-200), indicating that increasing the attention weights through chromatin loops enhances the model’s capacity to recognize distal cCRE-gene interactions.

**Fig.5.**
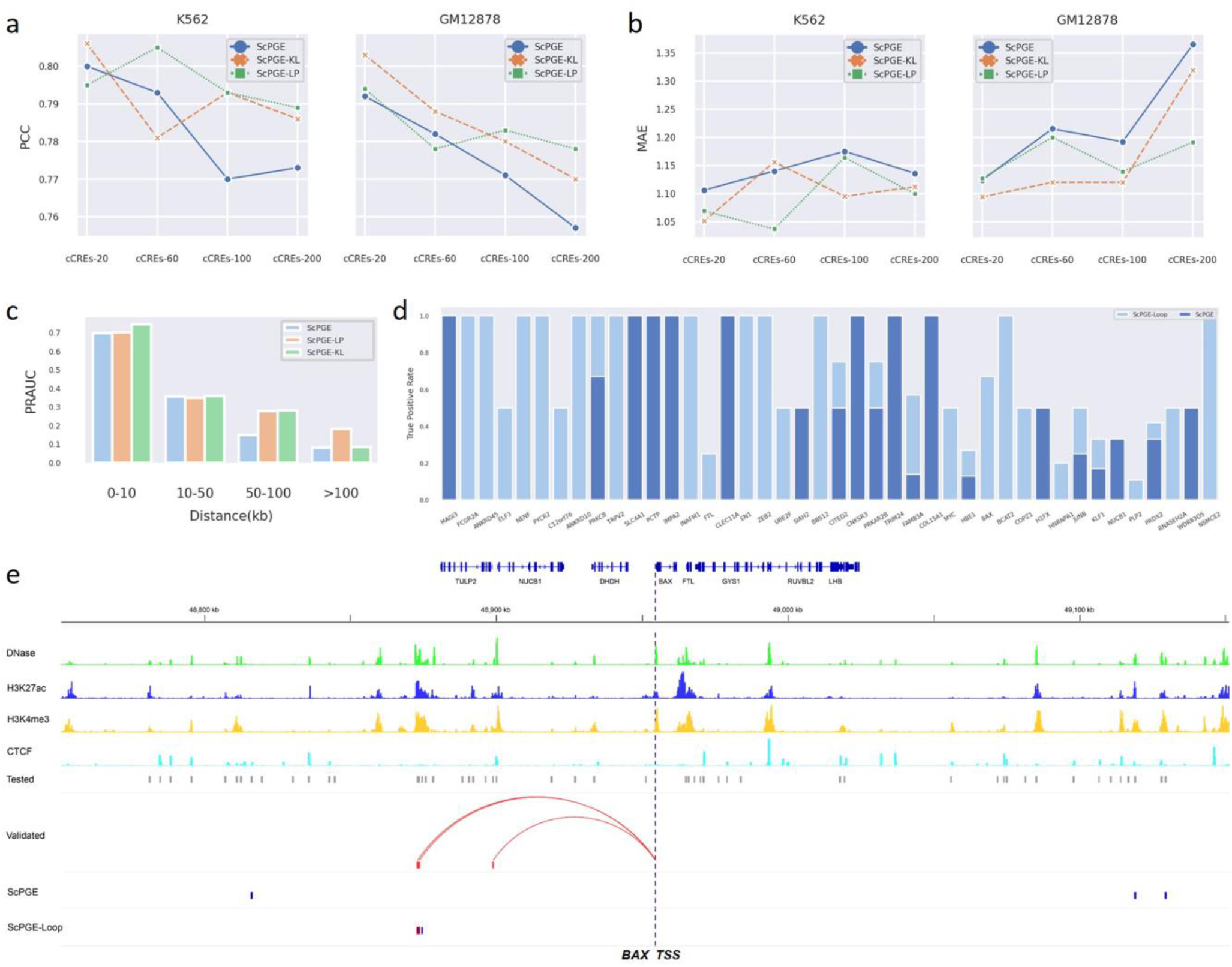
Chromatin loops facilitate ScPGE to capture cCRE-gene interactions. (a-b) Comparison of the performance of ScPGE, ScPGE-KL, and ScPGE-LP as the number of cCREs increases. (c) The performance (PRAUC) of ScPGE, ScPGE-KL, and ScPGE-LP in classifying cCRE-gene pairs at different distance groups. (d) The true positive rates of ScPGE and ScPGE-Loop on validated ccREs. (e) Visualization of active cCREs of the BAX gene missed by ScPGE but identified by ScPGE-Loop, where validated cCREs are represented by red squares and correctly identified cCREs are represented by red boxes.

Furthermore, we applied ScPGE-LP and ScPGE-KL to distinguish active enhancers from inactive enhancers in the K562-specific gene-enhancer interactions. As shown in Fig.5c, we find that both ScPGE-LP and ScPGE-KL outperform ScPGE (attention) at different distances. Moreover, ScPGE-LP performs better than ScPGE-KL at a longer distance (>100kb). These results further demonstrate the effectiveness of the two methods in identifying distal gene-enhancer interactions. Given that ScPGE-LP is proficient in recognizing distal interactions and ScPGE-KL is proficient in recognizing proximal interactions, we combined the two methods to identify validated enhancers of target genes. To be specific, ScPGE-KL was used to identify active enhancers within 100kb, and ScPGE-LP was used to identify active enhancers beyond 100kb. For simplicity, we refer to it as ScPGE-Loop. As before, we prioritized all enhancers and selected top-*k* enhancers, where *k* is the number of validated enhancers, to calculate the true positive rate (TPR). As shown in Fig.5d, ScPGE-Loop performs better than ScPGE on 30 out of 42 genes according to the TRP metric. For example, there are three validated enhancers (red boxes) surrounding the BAX gene. As a result, ScPGE (attention) fails to identify any validated enhancers, but ScPGE-Loop successfully identifies two of the three validated enhancers (Fig.5e). Taking the MYC gene as another example, ScPGE (attention) fails to identify any validated enhancers, but ScPGE-Loop successfully identifies one of the two validated enhancers (Supplementary Fig.8).

### 2.6 ScPGE captures important TF motifs

To capture cell type-specific TF motifs, we first selected all true positives (TPs) from the test set, and then ran the MEME-ChIP program(Machanick and Bailey 2011), which integrates multiple tools to perform comprehensive motif analysis on DNA sequences, to discover TF motifs from these TPs, and the most relevant discovered motifs were matched against the HOCOMOCO V11(Kulakovskiy et al. 2018) using TOMTOM(Gupta et al. 2007). As shown in Fig.6a, we identified a series of important TF motifs that are grouped by TF family and are often enriched in proximal or distal regulatory regions. For instance, SP1-like TFs activate or repress basal transcription by usually binding to the GC box (GGGCGG) or GT/CACC box in the promoter region of many genes. CTCF-like TFs not only function as transcriptional activators or repressors by binding to proximal or distal cCREs, but also block communication between enhancers and upstream promoters by binding to a transcriptional insulator element, thereby mediating the formation of three-dimensional structures of chromatin. Kr-like TFs, hematopoietic-specific transcription factors, could induce high-level expression of adult beta-globin and other erythroid genes by binding to the DNA sequence CCACACCCT in the beta hemoglobin promoter.

**Fig.6.**
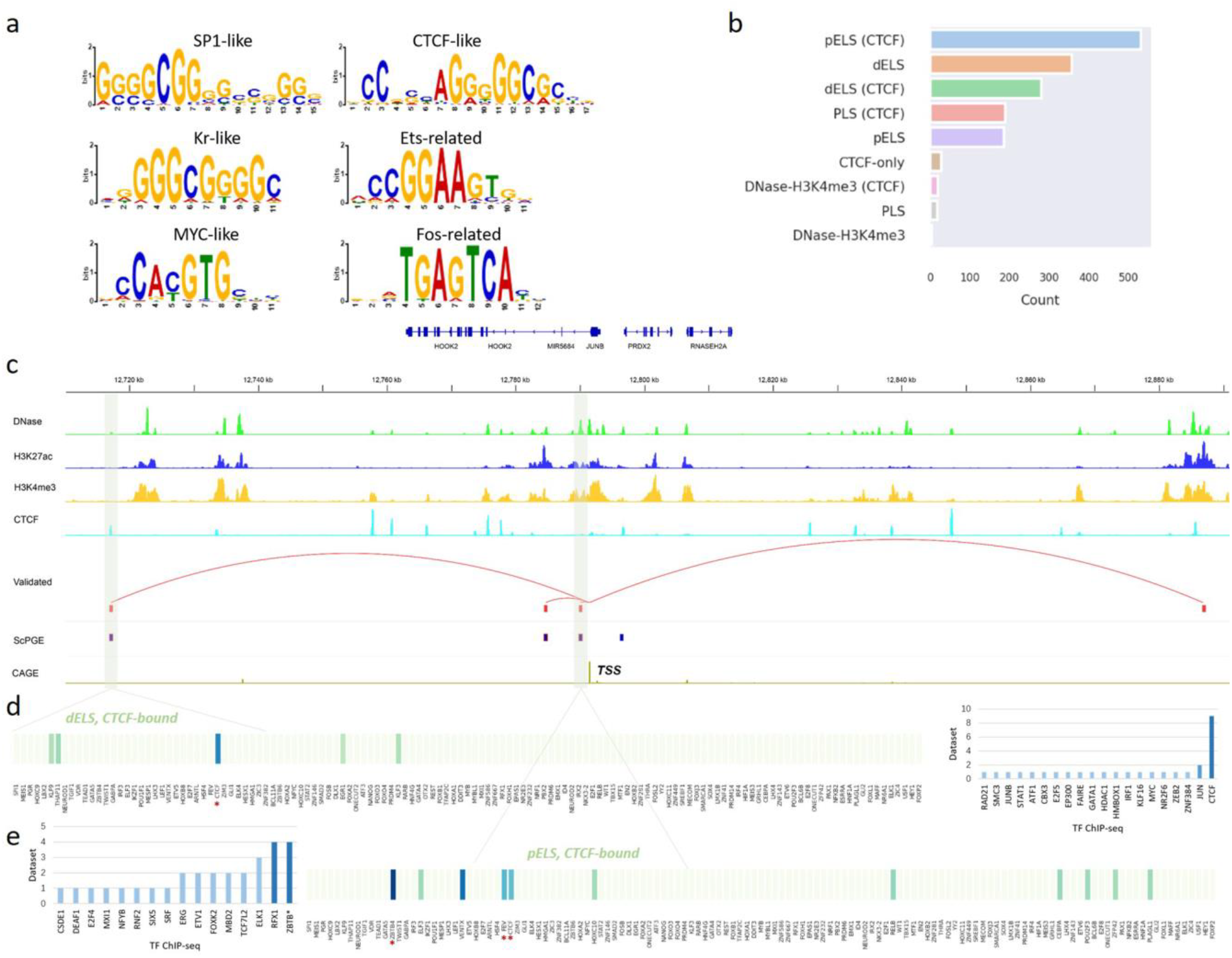
ScPGE discovers important motifs in specific types of cCREs. (a) Some important TF motifs found in true positives categorized by ScPGE, which are labelled by TF family names. (b) The distribution of types of cCREs, sourced from the SCREEN Registry V3, where *pELS* means proximal enhancer-like signatures, *dELS* means distal enhancer-like signatures, and *PLS* means promoter-like signatures. (c) Visualization of active cCREs of the JUNB gene identified by ScPGE, where validated cCREs are represented by red squares and correctly identified cCREs are represented by red boxes. (d) The contributions of TF motifs in a type-specific cCRE (dELS, CTCF-bound), where significant TFs are supported by TF ChIP-seq datasets. (e) The contributions of TF motifs in a type-specific cCRE (pELS, CTCF-bound), where significant TFs are supported by TF ChIP-seq datasets.

To further analyze the different functions of cCREs, we counted the distribution of types of active cCREs. Specifically, (i) we selected all TPs from the test set and computed the attention weights of all TPs using the ScPGE model; (ii) we extracted the middle row of the averaged attention matrix corresponding to the gene index and normalized these weights to a range of 0 to 1. Then, we sorted the weights and selected top-*k* (*k*=5) cCREs as the candidate set, denoted by ∅_a_; (iii) we performed *in-silico* perturbation using the ScPGE model and calculated the perturbation scores of all TPs. Then, we sorted these scores and selected top-*k* (*k*=5) cCREs as another candidate set, denoted by ∅_*p*_; (iv) we took the intersection of these two sets as the final active cCREs by ∅_a_ ∩ ∅_*p*_; (v) we counted the distribution of types of the final active cCREs. As shown in Fig.6b, we find that the most frequently regulated types of cCREs are proximal elements (e.g., pELS, PLS), followed by distal elements (e.g., dELS), and that most of these elements involve CTCF binding. This implies that CTCF is an important multifunctional transcription factor that is extensively involved in gene transcriptional regulatory activities.

For the JUNB gene, we picked out two cCREs that were correctly recognized by ScPGE from the four validated cCREs, one with the type dELS, CTCF-bound and another with the type pELS, CTCF-bound. For the distal cCRE, we calculated the contributions of TF binding scores by DeepLIFT(Shrikumar et al. 2017) and highlighted a few top TFs by ranking their contributions. As shown in Figs.6c-d, we observe that CTCF is a primary contributor, just matching the function of the cCRE labelled as dELS, CTCF-bound. Moreover, we searched all K562-specific ChIP-seq binding datasets in CistromeDB(Zheng et al. 2019) and found that the vast majority of CTCF ChIP-seq data had peaks falling in distal cCREs. For the proximal cCRE, the highlighted TFs are a group of proximal regulators, such as ZBTB4, VENTX, FEV (ETS family), CEBPA, POU2F3, which are partly supported by relevant ChIP-seq binding datasets. It has been reported in the literature that ZBTB4 is involved in the negative regulation of transcription by binding to RNA polymerase II, and VENTX may function as a transcriptional repressor, potentially playing a role in the maintenance of hemopoietic stem cells. Notably, although K562-specific CTCF ChIP-seq peaks are not enriched in the proximal cCRE, CTCF is still recognized as one of the important contributors, just matching the function of the cCRE labelled as pELS, CTCF-bound.

## 3. Discussion

Deciphering the relationships between cis-regulatory elements (CREs) and target gene expression has been a long-standing unsolved problem in molecular biology. To address this problem, some quantitative models have been proposed recently for modelling the relationships between cCREs and gene expression. For example, Enformer proposed a large transformer-based model to predict gene expression by taking in long-range genomic regions containing a sufficient number of cCREs. However, training such a large model with long-range sequences requires significant computing resources, posing a substantial challenge for researchers with limited computing resources. To mitigate this challenge, we proposed a scalable computational framework to predict gene expression by integrating DNA sequences, TF binding scores, and epigenomic tracks from discrete cCREs. A series of comparative experiments demonstrates the superiority and scalability of ScPGE. By comprehensively analyzing ScPGE’s predictions, we identified distinct regulatory patterns in TPs, FPs, TNs, and FNs, particularly a pattern in TPs where the regulatory effect of cCREs on genes decreases with distance. With this pattern, ScPGE can perform cross-cell or cross-species gene expression prediction with high accuracy, suggesting its potential application to previously unseen cell types or species. Additionally, motivated by this pattern, we developed two methods to improve the performance of identifying distal cCRE-gene interactions by incorporating chromatin loops into the ScPGE model.

By analyzing the contributions of TF motifs, we observed a phenomenon that a few important motifs belong to the same TF family and share a consensus binding sequence, which suggests a possible redundancy in the list of TF motifs. Compact and representative TF motifs through motif clustering analysis, therefore, may help further improve the performance of ScPGE. For example, the SCENIC+ motif collection includes 34,524 unique motifs gathered from 29 motif collections, which were clustered with a two-step strategy(Bravo González - Blas et al. 2023). The collection spans a total of 1,553 TFs, 1,357 TFs, and 467 TFs, respectively in human, mouse, and fly, providing comprehensive, compact, and representative TF motifs for different species. In addition, through analysis of predictions, the patterns found in FPs and FNs were similar to those in TPs and TNs, which may lead to incorrect predictions. Therefore, by utilizing these patterns, it is possible to exclude these noisy samples in advance, thereby further improving the performance of ScPGE. For example, if some samples in TNs satisfy the pattern found in TPs, these samples are considered noise samples and excluded; conversely, if some samples in TPs satisfy the pattern found in TNs, these samples are considered noise samples and excluded.

ScPGE stands out from gene expression prediction methods due to its lightweight design and flexible scalability. However, we found that the predictive performance of ScPGE decreases as the number of cCREs increases, implying that ScPGE struggles to capture the interactions between genes and distal cCREs effectively. Although chromatin loops were used to enhance the ability of ScPGE to capture the interactions between genes and distal cCREs, the predictive performance of ScPGE using distal cCREs remains relatively low. Given the powerful representational capabilities of large models(Dalla-Torre et al. 2025), we will leverage pre-trained DNA models and optimize a student model through knowledge distillation, which not only yields a lightweight model but also preserves the feature learning capabilities of large models. By doing so, we can mitigate to some extent the shortcoming that the model cannot effectively capture distal cCRE-gene interactions.

## 4. Methods

### Data preparation

#### Candidate Cis-regulatory elements (cCREs)

cCREs for Human were downloaded from the SCREEN Registry Version 3 (https://screen.encodeproject.org/). The Registry contains 1,063,878 human cCREs in GRCh38, which are derived from ENCODE data using four types of data including DNase, H3K4me3, H3K27ac, and CTCF signals. cCREs with DNase-only and low-DNase marks were removed. Given that CREs usually cluster together to form Cis-regulatory modules (CRMs)(Hardison and Taylor 2012), adjacent cCREs were merged within a window of size 600 bp, reducing the number of cCREs to 726,796. All cCREs were expanded to 600bp for the convenience of modelling.

#### Gene expression data

For RNA-seq, total RNA-seq and polyA plus RNA-seq data in 19 human cell types were downloaded from ENCODE. Released transcript quantifications mapped to GRCh38 and annotated to GENCODE V29 were retained. RNA-seq gene expression was calculated as the sum of all transcript’s TPM, and scaled by the log_10_(1+*x*) function.

For CAGE-seq, 10 in 19 human cell types, signals of all reads mapped to GRCh38 and annotated to GENCODE V24 were downloaded from ENCODE. CAGE-seq gene expression was determined by aggregating read signals within a 384-bp window centered on each gene’s unique TSS, following Enformer’s protocol, and scaled by the log_10_(1+*x*) function.

#### DNase-seq and ChIP-seq data

For each cell type, DNase-seq and ChIP-seq files mapped to GRCh38, including DNase, H3K4me3, H3K27ac, and CTCF types, were downloaded from ENCODE. Multiple read-depth normalized signal files for DNase-seq and fold change over control files for ChIP-seq were retained. Then, the signal files for the same type were merged using the *bigWigMerge* software from UCSC.

#### Active TF motifs

A full list of 769 TF motifs, represented by position frequency matrix (PFM), was downloaded from the HOCOMOCO V11(Kulakovskiy et al. 2018). Active TFs were defined as those with higher expression levels in a specific cell type. Therefore, all TFs were first sorted by their expression levels, and top 600 TFs in the full list were then selected as active TFs for a specific cell type. Finally, the active TF motifs were used to calculate cell-type-specific TF binding scores.

### Model design

#### Data construction

To incorporate more cCREs, Enformer took long-range continuous DNA sequences as input and dealt with them using *Conv1D* operations. However, cCREs are discretely distributed on both sides of genes and therefore cannot be directly handled by *Conv1D* operations. Inspired by our previous works(Zhang et al. 2018; Zhang et al. 2019), we assembled cCREs together with sequence features, TF binding scores, and epigenomic signals into 3-dimensional tensors. As shown in Supplementary Fig.1, the details of data construction are as follows:

##### Sequence features

DNA sequences of a gene and its surrounding cCREs were expanded to 600bp and transformed into one-hot matrices of shape (4×600) following the protocol {A: [1, 0, 0, 0]; C: [0, 1, 0, 0]; G: [0, 0, 1, 0]; T: [0, 0, 0, T]}. Then, one-hot matrices were separately reshaped into 4×1× 600 and assembled together along the second axis, forming a 3-dimensional tensor of shape 4×(*m*+1)×600 where *m* represents the number of cCREs and is a hyper-parameter with a default value of 20. After this processing, we can easily apply *Conv2D* operations to deal with these discrete cCREs simultaneously.

##### Epigenomic signals

According to the coordinates of cCREs on the genome, epigenomic signals (DNase, H3K4me3, H3K27ac, and CTCF) were extracted from their corresponding epigenomic files using the *pyBigWig* software and scaled by the log_10_(1+*x*) function. Following the same idea, the epigenomic signals of cCREs were reshaped and organized into a 3-dimensional tensor of shape 4×(*m*+1)×600 where ‘4’ denotes four types of data, and this parameter can be dynamically adjusted according to the availability of epigenomic data.

##### TF binding scores

A set of 600 active TF motifs was utilized to calculate the TF binding scores for cCREs. Specifically, for each cCRE, a motif represented by a position frequency matrix (PFM) was multiplied and summed with one-hot matrix of the cCRE, producing a vector. Then, the maximum value of the vector was taken as the TF binding score for this motif. As a result, this process generated a vector of 600 TF binding scores, which were subsequently normalized to a range of 0 to 1. Following the same idea, the TF binding scores of cCREs were reshaped and organized into a 3-dimensional tensor of shape 1×(*m*+1)×600.

##### Chromatin loops

Cell-type-specific chromatin loops at the 5kb resolution were downloaded from Loop Catalog(Reyna et al. 2025), such as K562 or GM12878, which were derived from H3k27ac HiChIP data by using the FitHiChIP utility(Bhattacharyya et al. 2019). If there exist interactions between cCREs and cCREs, as well as between genes and cCREs in chromatin loops, the normalized counts of chromatin loops were calculated and scaled by the log_10_(1+*x*) function, otherwise 0. As a result, for each gene and its surrounding cCREs, an interaction matrix of shape (*m*+1)×(*m*+1) was constructed.

#### Model architecture

As shown in Supplementary Fig.2, the backbone of ScPGE is composed of three key modules: (i) a feature-learning module to learn features of cCREs, and (ii) an interaction-learning module to model the relationships between genes and cCREs, as well as cCREs and cCREs, and (iii) a prediction module for predicting gene expression.

The feature-learning module is composed of 3 stem blocks for taking sequence features, TF binding scores, and epigenomic signals as input respectively, and 4 residual convolutional blocks for learning and integrating the three inputs. In the computational blocks, *Conv2D* and *MaxPool2D* are applied to deal with genes and discrete cCREs in parallel by setting kernel size to 1 along the height axis. At last, the feature-learning module uses a projection block to produce a feature embedding of fixed shape, e.g., (*m*+1)×256.

The interaction-learning module is composed of 4 transformer-based blocks for modelling the relationships between genes and cCREs. Each transformer block consists of a multi-head self-attention layer and a position-wise feed-forward network. In the self-attention layer, scaled dot-product attentions are performed as follows: the query 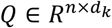, key 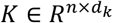, and value 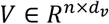 are calculated through a linear projection where *n* denotes the number of embeddings and *d*_*k*_, *d*_*v*_ the number of channels; the attention weights are calculated by 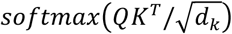 representing the attention pairwise; the value representing the semantics of all embeddings is aggregated according to the attention weights. For position embedding, we follow T5(Raffel et al. 2020) to apply a relative positional embedding 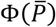 onto the attention weights, where 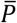 is the relative position between a gene and its cCREs. Additionally, we mask the attention weights between genes to encourage the model to concentrate on capturing the relationships between genes and cCREs. The feed-forward network integrates the outputs from multi-head attention layers and introduces non-linearity. The calculation process can be described as the following equitation:

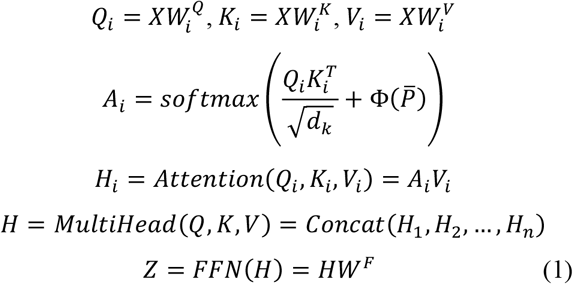

Where 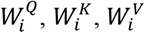 represent the weights of three linear layers respectively, and *H*_*i*_ denotes the outputs from the *i*-th self-attention layer, and *W*^*F*^ denotes the weights of the feed-forward network.

The prediction module is composed of a fully-connected feed-forward network and a soft plus activation to predict the gene expression. Through the last transformer block, a matrix *Z* of shape (*m*+1)×*d* is generated where *d* denotes the output dimension of the last transformer block, e.g., 256. To encourage the model to concentrate on gene expression prediction, the middle row of *Z* corresponding to the gene index is extracted, representing the relationships between a gene and its cCREs. Then, the prediction module takes the middle row as input and predict the expression level of the gene.

#### Model training

Since gene expression prediction is a regression task, the Mean Squared Error (MSE) was employed to calculate the differences between true and predicted expression levels (Equation 2).

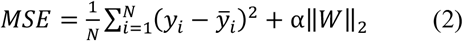

where *N* is the number of samples, *y*_*i*_ and 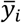 are the true and predicted expression levels, respectively; α is a regularization parameter to leverage the trade-off between the fitting and generalizability of ScPGE; || ||_2_ indicates the L2 norm.

Given that deep learning-based models are sensitive to parameter initialization, we adopted a ‘warm-up’ strategy to reduce the impact of parameter initialization. Specifically, we first warmed up ScPGE by running a few randomly-initialized models and selecting the best-performing model in terms of validation performance. Then, we resumed to train the best-performing model with a batch size of 64 using the MSE loss and AdamW with default hyper-parameters, except that the initial learning rate was gradually increased from 1e-06 to 1e-03 during the first stage (5000 steps) and decreased from 1e-03 to 1e-06 during the second stage (3000 steps). The model was validated every 1000 steps and the best-performing model is stored. All ScPGE models were implemented in PyTorch and trained on one A100 GPU.

#### Incorporation of chromatin loops

With the advent of technologies such as Hi-C and HiChIP for genome-wide chromatin interaction measurements, a large amount of chromatin interactions has been generated in a variety of cellular environments. Apparently, these interactions can provide highly useful information for guiding ScPGE to capture the relationships between genes and cCREs, but this information has been less utilized by current state-of-the-art models. To incorporate chromatin loops into the ScPGE model, we developed two methods: a direct method and an indirect method.

In the direct method, we directly put the interaction matrix *M*, representing cell-type-specific chromatin loops, into the self-attention layer with the purpose of increasing the attention weights of physical chromatin interactions. Through this method, the ScPGE model was guided to pay more attention to physical CRE-gene interactions. Correspondingly, the equation for computing *A*_*i*_ is modified as follows:

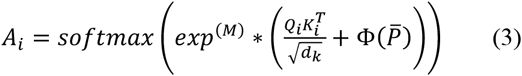

In the indirect method, we added a new loss to the MSE loss, which is defined as a KL Divergence loss between the interaction matrix and attention weights, with the goal of aligning their distributions by training a joint loss (Equation 4). Specifically, (i) the multi-head attention weights were extracted from transformer blocks and subsequently averaged element by element across all heads and layers, denoted by *Ā*; (ii) the middle rows of *Ā* and *M* corresponding to the gene index were extracted, represented by *Ā*_*g*_ and *M*_*g*_ respectively; (iii) given that *M*_*g*_ is extremely sparse, even with all values being 0, and inspired by the pattern found in this study that the regulatory effect of cCREs on target genes would decrease with distance, we added a general regulatory effector *R*_*g*_ to *M*_*g*_, which can be simulated with a function *exp*^−|*x*|/2^; (iv) after that, *M*_*g*_ and *Ā*_*g*_ were normalized to a range of 0 to 1.

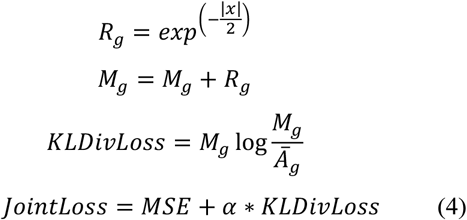

where *x* denotes the distances between a gene and its surrounding cCREs, and *α* is a hyper-parameter that specifies the contribution of the two losses to the prediction. We set it to 1 in this study.

### Model interpretability

#### Attention weights

The multi-head attention weights were extracted from all transformer blocks and subsequently averaged element-wise across all heads and layers. Since we focused on the attentions from gene to cCREs, the middle row of the averaged attention matrix corresponding to the gene index was extracted and then normalized to a range of 0 to 1, reflecting the relationships between the target gene and its surrounding cCREs. Therefore, we can directly investigate the importance of each cCRE to the target gene by matching the query index at cCREs.

#### In-silico perturbation

ScPGE predicts gene expression levels by utilizing the multimodal information of cCREs. Therefore, we can investigate the effect of cCREs on gene expression prediction by perturbing each cCRE. *In-silico* perturbation was performed by calculating the gene expression changes before and after masking each cCRE, |*G*_*o*_ − *G*_*p*_|/*G*_*o*_.

#### Contribution

DeepLIFT(Shrikumar et al. 2017), a feature attribution method for computing the contribution of each feature in an input to a scalar prediction from a neural network model, was utilized to calculate the contributions of TF binding scores. Specifically, (i) the contributions of TF binding scores were computed by DeepLIFT, generating a matrix with a shape of 1×(*m*+1)×600; (ii) after selecting the target cCRE, a vector with a shape of 1×600 was generated, where ‘600’ represents the number of TF motifs, and then TF motifs were ranked by their corresponding contributions; (iii) top-*k* (e.g., *k*=10) TF motifs were selected as the highlighted ones. This process was implemented via Captum (v0.6.0)(Kokhlikyan et al. 2020), which is a PyTorch platform for model interpretability.

### Categorization of predictions

To discover different patterns of false positives (FPs), true positives (TPs), false negatives (FNs), and true negatives (TNs), the predictions from the test set were categorized into FPs, TPs, FNs, and TNs by following the two steps: (i) the predicted and true gene expression levels were converted to a range of 0 to 1 by the equation: (*x* − *x*_*min*_)/(*x*_*max*_ − *x*_*min*_); (ii) we defined them as FPs by the rule: the prediction (*P*) is above 0.7 and the true (*T*) is below 0.2, as TPs by the rule: both *P* and *T* are above 0.7, as FNs by the rule: *P* is above 0.2 and *T* is below 0.7, and as TNs by the rule: both *P* and *T* are below 0.2.

### Cross-species gene expression prediction

For the GM12878 and K562 cell lines from Human, we selected the CH12.LX and MEL cell lines from Mouse as targets for cross-species prediction, respectively. For CH12.LX and MEL, we downloaded cCREs for Mouse from the SCREEN Registry V3, total RNA-seq and polyA plus RNA-seq data annotated to GENCODE vM21, DNase-seq and ChIP-seq files mapped to GRCm38, as well as active TF motifs for Mouse from the HOCOMOCO V11, and followed the process of data preparation described above to prepare all relevant data. For each cell type, all coding genes on chromosome 16 were used for validation, all coding genes on chromosomes 8 and 9 were used for testing, and all coding genes on the remaining chromosomes, except chromosome Y, were used for training. After that, we trained the ScPGE models for CH12.LX and MEL using the corresponding training data.

### Baseline methods

Three state-of-the-art models, including Enformer(Avsec et al. 2021), CREaTor(Li et al. 2023), and EPInformer(Lin et al. 2024), serve as the baseline methods for gene expression prediction. Enformer, a deep neural network that integrates CNN and transformer, takes long-range DNA sequences of length 196 kbp as input to predict multiple genomic tracks of human and mouse genomes simultaneously. CREaTor, a hierarchical attention-based deep neural network, utilizes cCREs in open chromatin regions together with ChIP-seq of transcription factors and histone modifications to predict the expression level of target genes. CREaTor can model cell-type-specific cis-regulatory patterns in new cell types without prior knowledge or additional training. EPInformer introduces a transformer-based framework to improve gene expression prediction by integrating promoter-enhancer interactions with their sequences, epigenomic signals, and chromatin contacts.

For comparing with Enformer, pre-trained Enformer models were downloaded from https://github.com/google-deepmind/deepmind-research/tree/master/enformer, and genomic sequences flanking genes of interest were prepared following the original study’s instructions. For a specific cell type, all predictions matching that cell type were extracted and aggregated, and then the gene expression level was determined by aggregating the predicted signals within a 384-bp window centered on each gene’s TSS. To ensure a fair comparison, the ScPGE models were trained using the multimodal information of cCREs to predict CAGE-seq gene expression.

Following the instructions for running CREaTor, we retrained the CREaTor models using DNA sequences and four types of epigenomic signals across 19 human cell types with default hyper-parameter settings. Similarly, following the guidelines for training EPInformer from scratch, we retrained the EPInformer models using the original inputs, except for chromatin contacts, across 19 human cell types with default hyper-parameter settings, since chromatin contacts are not available for most cell types.

### cCRE-gene interactions

Three K562-specific enhancer-gene interaction datasets were collected from Fulco *et al*.(Fulco et al. 2019), Gasperini *et al*.(Gasperini et al. 2019), and Schraivogel *et al*.(Schraivogel et al. 2020), in which enhancer-gene pairs were categorized into four groups based on distance, including ‘<10kb’, ‘10-50kb’, ‘50-100kb’, and ‘>100kb’ groups. For all three datasets, the genomic coordinates of all candidate enhancers were first converted from hg19 to hg38 using UCSC’s liftover software and then integrated together by genes.

## Data Availability

RNA expression, DNase-seq, ChIP-seq, and CAGE files were downloaded from https://www.encodeproject.org/ (Supplementary Tables 1-3). TF motifs (PFM) were downloaded from the HOCOMOCO V11 (https://hocomoco11.autosome.org/). Cell-type-specific chromatin loops at the 5kb resolution were obtained from https://loopcatalog.lji.org/ (Supplementary Tables 4-5). cCREs for Human were downloaded from the SCREEN Registry Version 3 (https://screen.encodeproject.org/). CRISPR perturbation experiments of cCRE-gene interactions were collected from CREaTor (Supplementary Table 6).

## Code Availability

The ScPGE algorithm was implemented in PyTorch. The source code is available at https://github.com/turningpoint1988/ScPGE.

## Acknowledgments

This work was supported in part by the National Natural Science Foundation of China, Nos. 62372255, 62333018, W2412087, 62402250, 62432013, 62433001, and partly supported by the Natural Science Foundation of Zhejiang Province, No.LMS25F020001, and supported by the Key Research and Development Program of Ningbo City under Grant Nos. 2024Z112, 2023Z219, 2023Z226, and supported by the Yongjiang Talent Project of Ningbo, Yongrencaifa No.2024-4, and supported by the Basic Research Program Project of Department of Science and Technology of Guizhou Province, No.ZK2024ZD035, and supported by the Youth Innovation Team of Colleges and Universities in Shandong Province (2023KJ329), and supported by the IDT High Performance Computing Platform for providing computational resources for this project.

## Author Contributions

D. Huang and Q. Zhang conceived the basic idea; Q. Zhang designed experiments, developed algorithms, and wrote the manuscript; S. Wang and Z. Li carried out the data analysis and model interpretation; W. Bao and W. Liu provided some suggestions for writing the manuscript and designing the experiments.

## Conflict of Interest

The authors declare no conflict of interest.

## Notes

### Competing Interest Statement

The authors have declared no competing interest.

